# Case-specific selection of batch correction methods for integrating single-cell transcriptomic data from different sources

**DOI:** 10.1101/2024.05.26.595911

**Authors:** Xiaoyue Hu, He Li, Ming Chen, Junbin Qian, Hangjin Jiang

**Author notes:** These authors contribute equally to this work (Xiaoyue Hu, He Li).

## Abstract

Integrating single-cell RNA-sequencing datasets from different sources is a common practice to empower in-depth interrogation for biological insights, where batch effect correction (BEC) is of vital importance. However, an inappropriate BEC may lead to overcorrection and report misleading results on downstream analyses including cell annotation, trajectory inference and cell-cell communication. Hence, we develop the Reference-based Batch Effect Testing (RBET), a novel statistical framework for evaluating the performance of different BEC methods by leveraging housekeeping-gene inspired reference genes and MAC statistics for distribution comparison. Comparing with existing methods, RBET is more powerful on detecting batch effect, overcorrection sensitive, computationally efficient, and robust to large batch effect sizes. Furthermore, extensive multi-scenario real examples show that RBET selects optimal BEC tools for consistent downstream analysis results, which confirm prior biological knowledge. This comprehensive BEC decision-making tool is available as an R package.

## Introduction

Single-cell RNA sequencing (scRNA-seq) provides an unprecedented opportunity to define molecular cell type and cell state in a data-driven fashion, and has achieved great accomplishment in exploring the mechanism of different diseases at the cell level^1–3^. Researchers may take the advantage to investigate in-depth biological insights by integrating massive datasets, either publically available or generated in-house, for multiple downstream analyses, such as cell type and state annotation, differential gene expression, trajectory inference, cell-cell communication, etc. But the underlying difficulty is the large variations or batch effect in these datasets, which are collected from different labs, experiments, capturing times, handling personnel, and even technology platforms, etc.^4,5^. These differences confound the true biological variations during data integration, which need to be removed by Batch Effect Correction (BEC) tools to ensure biologically meaningful results^6,7^. Indeed, various BEC methods have been proposed in recent years (**Fig. 1A, 1B**), and a recent benchmark study compared 14 BEC methods in terms of batch mixing, accuracy of cell type annotation, computational efficiency for large datasets, etc., and recommended Harmony, LIGER, and Seurat as top performers^8^.

**Figure 1.**
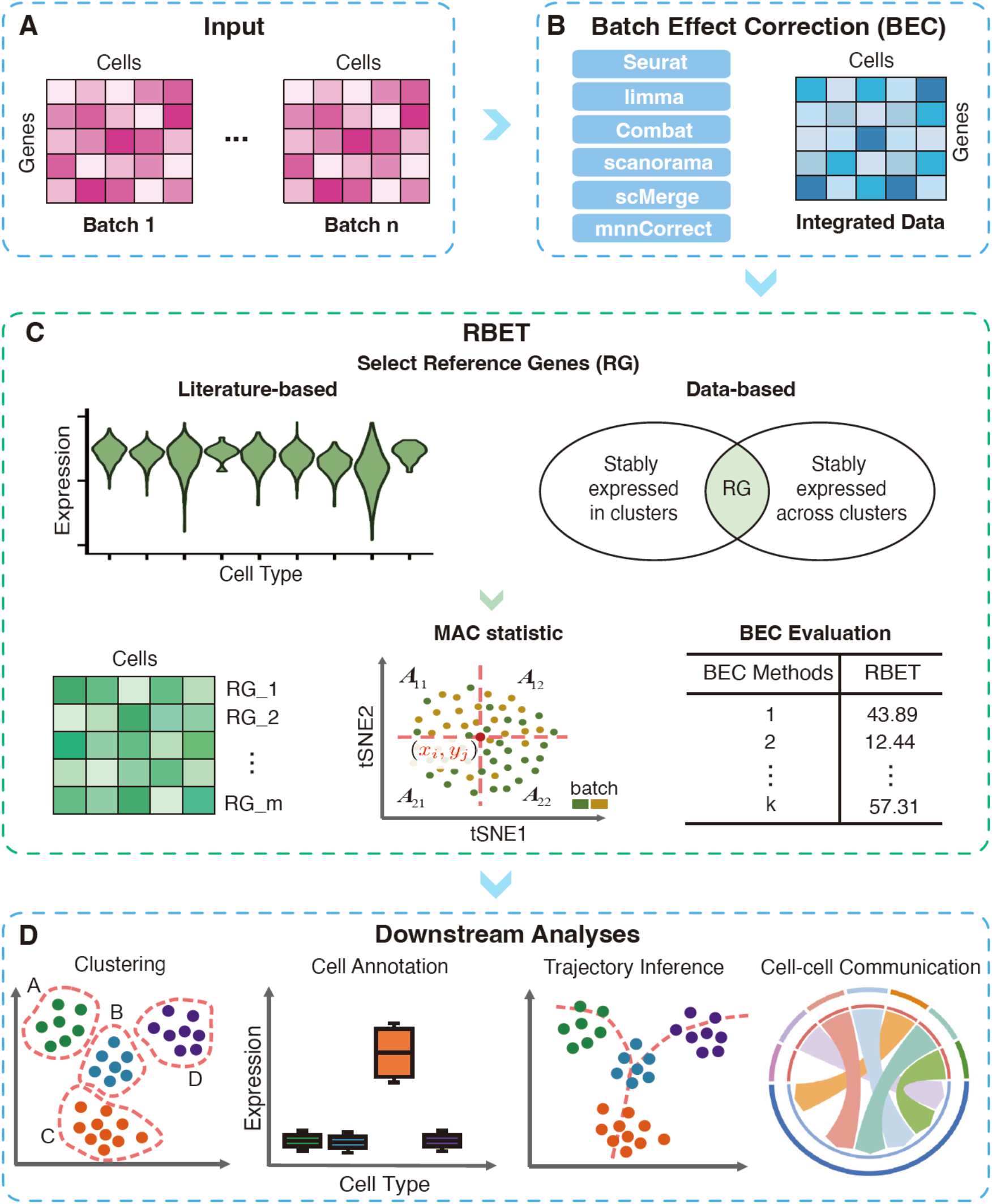
Overview of the workflow. **(A)** Single-cell gene expression matrices as input data from different sources. **(B)** Data integration performed using six different BEC tools. **(C)** RBET framework (See **Methods**). RBET selects the BEC method with smallest RBET value. **(D)** The performance of RBET is validated through downstream analyses including clustering, cell annotation, trajectory inference, and cell-cell communication.

However, these benchmarking studies lack comparisons for key metrics that could impact routinely performed downstream analyses such as trajectory inference and cell-cell communication. First, no difference should be detected between samples from different batches after BEC, both locally and globally. However, the existing BEC evaluation tools such as kBET^9^ or LISI^7^, may have a lower power in some cases where only parts of cells have batch effect. Second, these tools lack algorithm to estimate overcorrection, that is, degradation of true biological information while eliminating technical bias^9^. Overcorrection makes downstream analysis highly problematic that often leads to wrong conclusion. Third, BECs should be evaluated for capability to cope with large batch effect size. Previous benchmarking studies ultilized datasets with limited batch numbers (2-5) and arguably not sufficient batch effect sizes, while these are possibly several order of magnitude higher in real practice, given the overwhelming availability of scRNA-seq datasets^10^. Thus, a comprehensive and reliable test metric taking full consideration of the aforementioned scenarios is an unmet need for BEC applications.

Here, we propose a novel statistical method, Reference-based Batch Effect Testing (RBET) for evaluating the performance of BEC tools. This method takes advantage of reference gene (RG) expression/variation pattern and maximum adjusted chi-squared (MAC) statistics for two-sample distribution comparison. In both simulated and real data, we demonstrated a superior performance of RBET in detecting batch effects with overcorrection awareness, large batch effect robustness and high computational efficiency, as compared to kBET and LISI. More importantly, RBET excels in prioritizing BEC tools that feed downstream analyses for results consistent with prior biological knowledge.

## Results

### Overview of RBET framework

Housekeeping genes usually refer to genes that are essential for basic cellular functions and the maintenance of cellular viability^11^, and show uniform expression across various cell types under diverse conditions. Given their consistency, they are often used as internal controls and standards for the expression normalization and quantification^12–14^. Indeed, since housekeeping genes are stably and abundantly expressed, thus almost unaffected by dropout events, which are common for most other genes in scRNA-seq data. This is important because the inflated zeros in gene expression comprise both biological and technical variation, which are hard to discern. When evaluating batch correction for a human peripheral blood mononuclear cell (PBMC) dataset by Seurat, we found that the expression of housekeeping genes (i.e., RPS17 and RPS18 in **Fig. S1A-S1B)** was changed not only in expression level, but also in expression variation within the cell types before and after BEC, suggesting a potential usage for separating technical and biological sources of variation. Indeed, a perfect batch correction should align the expression level of housekeeping genes between batches (technical batch correction), but not at the expense of losing expression variation within the batch, or in other word, loss of biological information that causes overcorrection.

Based on this hypothesis, we proposed the RBET framework for evaluating the performance of different BEC methods (**Fig. 1C**). RBET is comprised of two steps: (1) selecting reference genes specific to each tissue or dataset; and (2) detecting batch effect on these RGs in the integrated datasets. The performance of RBET is then validated through downstream analyses, including clustering and cell type annotation, trajectory inference, and cell-cell communication (**Fig. 1D**). We established two approaches to select appropriate reference genes. One curated existing validated tissue-specific housekeeping genes as RGs from published literatures (**Fig. 1C**, left).For cases where validated tissue-specific housekeeping genes are not available, we directly calculated the RGs from the datasets of interest, assuming that the tissue-specific RGs should be stably expressed both within and across phenotypically different clusters (**Fig. 1C**, right; details see **Methods**). These RGs were then fed into the batch effect test. Note that testing the batch effect between two samples equals to compare their underlying distributions in a high-dimensional setting. Previously, we designed MAC statistics^16^ for the later problem in low dimensional cases. Hence, we proposed to map the original datasets into a two-dimensional space, i.e., tSNE or UMAP, and use MAC statistics for batch effect detecting (see **Methods**). Considering the large scale of scRNA-seq datasets, a sub-sampling strategy was proposed to reduce the computational cost while keeping the theoretical property of MAC statistics for batch effect detection (see **Methods**).

To distinguish RBET with different RG strategies, we use RBET for RBET with literature-based RGs, and data-based RBET for RBET with data-based RGs. The result is easy to interpret: a lower RBET score value implies better batch correction, and weaker batch effect.

### RBET outperforms the state-of-the-art in simulations

We first compared the batch effect detection performance between RBET, kBET and LISI under two simulation strategies: (1) Gaussian examples with different means or covariance structures modeling different patterns of batch effect as a toy model; and (2) examples with simulated gene expression level mimicking real data under different settings of cell type numbers, batch effect sizes and the proportion of cells having batch effect (see **Methods)**. In our simulations, the power of each method was evaluated from 200 independent repetitions under significance level 0.05. Generally, a larger sub-sample k of RBET resulted in a higher power but also needs more computational cost. We set k = 50 because k ≥50 already showed a stable performance in both Gaussian examples (**Fig. S2A**) and simulated gene expression data (**Fig. 2A** and **Fig. S2B**).

**Figure 2:**
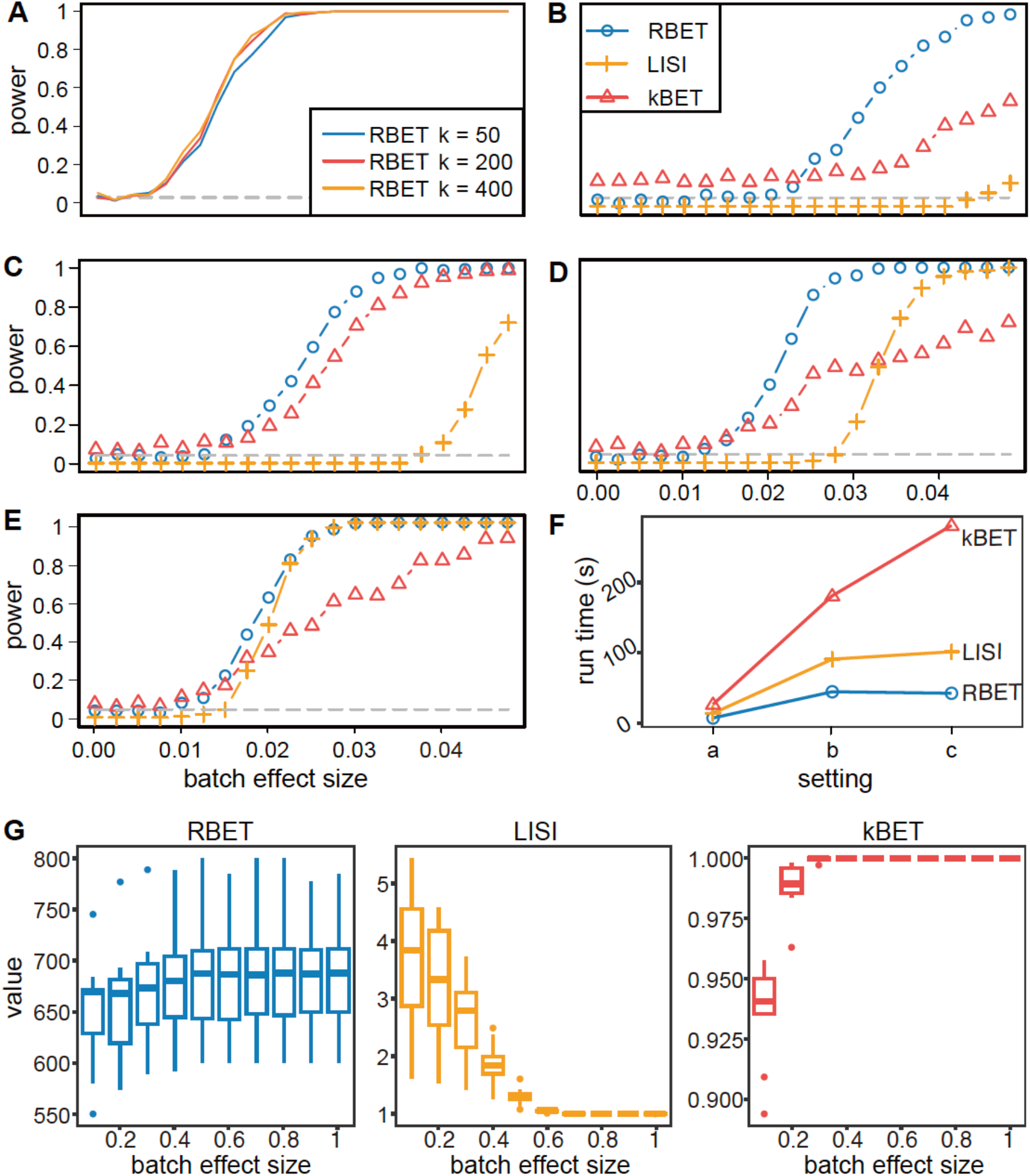
RBET outperforms other batch effect detection tools in simulated data. **(A)** The power of RBET under different sub-sampling size k in the simulated gene expression example with one cell type. The gray dotted line presents the theoretical power (0.05) of a reasonable method when null hypothesis is true, i.e., there is no batch effect. **(B-E)** The power of RBET, kBET and LISI under different batch effect sizes in the simulated gene expression example. Each batch contains two cell types and only one of them has batch effect. We consider the impact of the proportion of cells (*p*) having batch effect on the power of each method. **(B)** *p* = 20%. **(C)** *p* = 40%. **(D)** *p* = 60%. **(E)** *p* = 80%. **(F)** The total computational time used for 200 repetitions in the simulated gene expression example under three settings: (a) 1000 cells and 2000 genes in each batch; (b) 2000 cells and 4000 genes in each batch; (c) 4000 cells and 8000 genes in each batch. **(G)** The values of batch effect detection tools under different batch effect sizes.

In simple Gaussian examples, RBET showed comparable performance with kBET and LISI with different means or covariance structures (**Fig. S2C-S2E**). When turning to examples with simulated gene expression data, RBET outperformed other methods in terms of type I error control and a higher detection power, while LISI lost control over type I error (**Fig. S2F**). Importantly, we observed that the proportion of cells with batch effect, which is variable in real data scenario, exhibited a strong impact on the LISI performance (**Fig. S2G**). To challenge more difficult scenarios where only a subpopulation of cells had batch effect, we designed examples in which each batch had multiple cell types (2-8) but only parts of them had batch effect (see **Methods**). RBET always achieved the best performance in detection power in each case (**Fig. 2B-2E, Fig. S2H-S2I**), while kBET and LISI tended to lose control over type I error (**Fig. 2C, Fig. S2H-S2I**). Moreover, RBET also topped the computational efficiency test, demonstrating its potential to scale up for high batch number in big dataset (**Fig. 2F**). Since such scaled scenario may introduce bigger batch effect size, we tested RBET, kBET and LISI for their detection range by artificially increasing the batch effect size (**Fig. 2G**). Strikingly, RBET values remained variable across full batch effect size range, while the variations of LISI and kBET results started to collapse into a zero from the sizes of 0.5 and 0.2, respectively, indicative of reduced discrimination capacity on datasets with high batch effect. Overall, RBET outperformed the other two methods on multi-scenario simulated datasets in terms of detecting power, type I error control, computational efficiency and robustness to batch effect.

### RBET is aware of overcorrection

Change in highly expressed genes (HVGs) is often used to assess data overcorrection, but there is a lack of a consensus way to simulate overcorrection. We reason that some BEC algorithms, i.e., CCA method in Seurat V4, use nearest neighbour as foundation, and an increase on number of anchors k.anchor could potentially lead to overcorrection. Indeed, after increasing k.anchor in Seurat CCA function from 1 up to 100, we found that increasing k.anchor to 100 caused polarization of cells by batches as compared to 10 that showed well mixed cell distribution, suggesting loss of variance within the cluster (**Fig. 3A**). This was supported by a gradual decrease in the number of HVGs (**Fig. 3B**), as well as respective and total differentially expressed genes (DEGs, **Fig. 3C**) between cell groups detected in two batches after BEC. Both indicate a loss of true biological variation therefore an overcorrection. Further, although the RGs should be well aligned after BEC, their variation should be maintained, and thus their reduction may serve as another proxy metric to indicate overcorrection. Indeed, an increase on k.anchor led to loss of expression variation for RGs (**Fig. 3D**), confirming a successful simulation of overcorrection. In addition, this also highlights the importance of incorporating RGs in RBET algorithm. Accordingly, we found that RBET value decreased gradually in the first phase until k.anchor reached 10 (minimal value) (**Fig. 3E**), while a further increase of k.anchor led to gradual increase of RBET value at the second phase, which coincided with an increased level of overcorrection. Such biphasic change was not seen for kBET and LISI. Overall, these results suggest that RBET is overcorrection sensitive while LISI and kBET are not.

**Figure 3.**
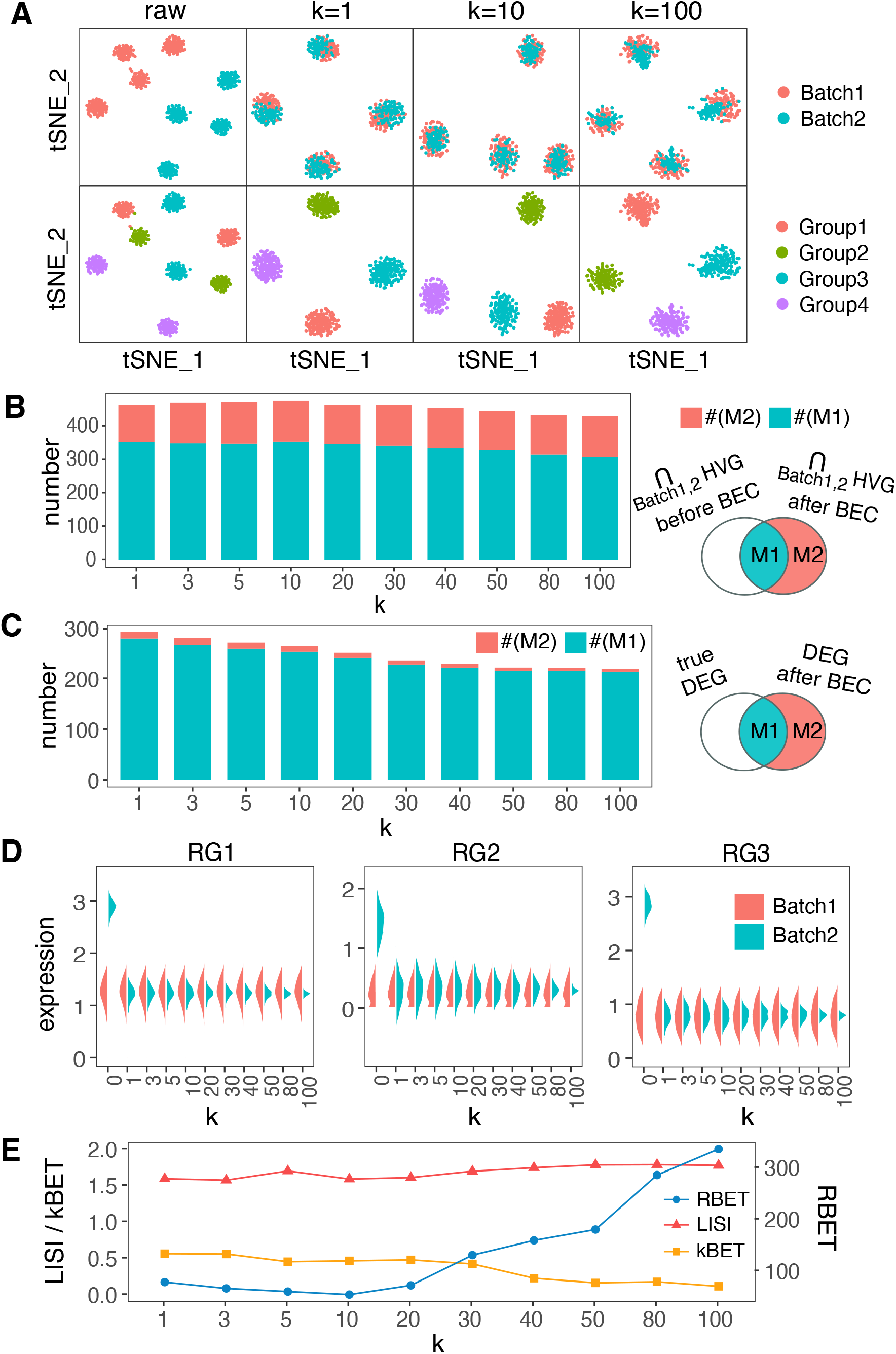
RBET detects batch effect and data overcorrection. We simulate 2 batches of cells, each with 4 groups. 100% of cells have batch effect, and BEC is performed by Seurat (v4.1.0) with different k.anchor (k). **(A)** tSNE plots show examples of simulated data before and after BEC. **(B)** The number of HVGs co-identified by two batches after BEC and their overlap with HVGs before BEC. **(C)** The number of identified DEGs after BEC and their overlap with true DEGs. **(D)** The expression patterns of reference genes (RGs) before and after BEC. **(E)** The values of RBET, kBET and LISI.

### RBET confirms results on cell type annotation

Since most scRNA-seq downstream analyses require full gene expression matrices, we focused on six BEC tools capable of returning full dimensional data, including Seurat^17^, scanorama^18^, scMerge^19^, limma^20^, Combat^21^ and mnnCorrect^22^ (**Fig. 1B**). Tools like Harmony^7^ and fastMNN^23^, which only output low-dimensional embedding, were not further explored in this study. To benchmark the power of RBET, we started from the pancreas dataset that included 3 technical batches (Celseq, Celseq2 and SMARTseq2) and 13 cell types with varied number of cells in each cell type (7∼2065)^24^ (**Fig. 4A**). To evaluate RBET performance, we curated experimentally validated housekeeping genes specific to pancreas as candidate RGs^25^, and checked whether they were differentially expressed across batches (**Table S1**, see **Methods**). Cells from different batches had strong batch effect, as evidenced by not well mixed clusters in UMAP (**Fig. 4B**), as well as large values of RBET and kBET, and a small value of LISI, respectively. Indeed, the smallest values obtained by RBET and kBET indicate the optimal BEC, whereas the largest LISI value signals a recommendation. After data integration by six BEC tools, both RBET and LISI selected Seurat as the best method, while kBET favored Combat (**Fig. 4C**). Data-based RBET also selects Seurat (**Table S5**), consistent with RBET (**Fig. 4C**). However, Combat clusters were apparently not well mixed by batches as visually inspected on UMAP (**Fig. 4C**), suggesting a low performance of kBET in this case.

**Figure 4.**
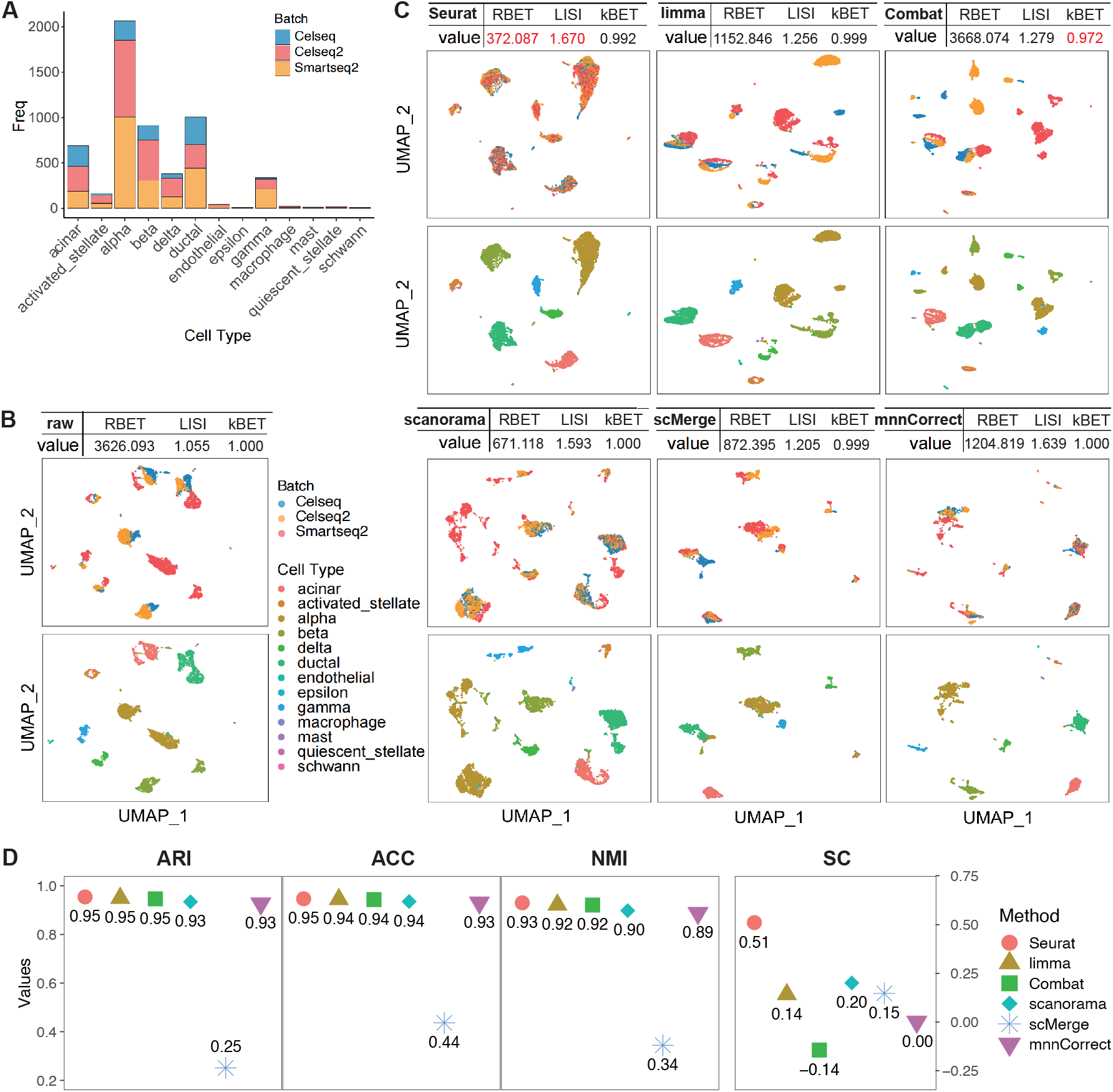
RBET selects valid BEC methods in pancreas data. **(A)** Histogram of cell type composition in each batch. **(B)** UMAP visualization of original dataset colored by batches (up) and cell types (down). **(C)** UMAP visualization of integrated data colored by batches (up) and cell types (down). The table in the title of each subfigure gives the BEC method used for integration and the evaluation of integration result given by RBET, LISI and kBET, whose optimal choices are colored in red. **(D)** The consistency (measured by ACC, ARI, NMI and SC) between real cell annotations and the annotations based on different BEC methods. Note that ACC, ARI, and NMI range in [0,1], while SC ranges in [-1,1], and a larger value indicates a better annotation result.

Next, we used cell annotation to further validate the selections. The cell types were annotated by ScType^26^ with the help of marker genes^27^. We compared cell annotation results from different BEC methods with real cell tags using accuracy (ACC), adjusted rand index (ARI), normalized mutual information (NMI) and silhouette coefficient (SC). For ACC, ARI and NMI, all six BEC tools obtained desired scores, expect for scMerge (**Fig. 4D**), which scored the lowest for all three metrics, although it had a clean cell clustering on UMAP. Indeed, when we looked into the marker gene expression patterns for the major cell types in the scMerge dataset, we observed evenly expressed profile across different cell types (**Fig. S4A**). In addition, a failure in RGs expression alignment and partial loss of RGs expression variation may also contribute to bad BEC results (**Fig. S4B**). Finally, Seurat obtained the highest SC score, confirming the correct choice by RBET and LISI. However, when choosing the second best, RBET’s scanorama beated LISI’s mnnCorrection in ACC, NMI and SC (**Fig. 4D**).

### RBET ensures accurate trajectory analysis

Trajectory analysis infers dynamic process of cell differentiation and state transition, and provides great insights into the biological phenomena^28,29^. In this section, we validated the performance of RBET for single-lineage monocyte^30^ and multi-lineage CD8^+^ T cell development^31,32^ with both literature-based RGs (**Table S2, S3**) and data-based RGs. The monocyte data contained 10,878 monocyte cells from the same healthy human blood, which were sampled and processed in three batches at different times (**Fig. 5A**). The batch effect was corrected by six BEC methods at various degrees, except for scMerge, which failed to incorporate batch3 (**Fig. 5B**). RBET, LISI and kBET had different optimal choices, including Combat, scanorama and mnnCorrect, respectively. Two versions of RBET gives the same selection (**Fig. 5B, Table S5**). kBET’s choice mnnCorrect appeared not ideal since their clusters were not well mixed for batches in UMAP as compared to Combat and scanorama. To nail down the ideal BEC tool for downstream trajectory analysis, we took advantage of a top rated tree-based tool for trajectory inference, Slingshot^28^. Consistent with monocyte biology, this analysis revealed a continuous single-ended trajectory from CD14^+^ monocytes (classical) to CD16^+^ monocytes (non-classical) by Seurat, limma, Combat and scanorama, while trajectories from scMerge and mnnCorrect were not biologically meaningful as they were either discontinuous or branched (**Fig. 5C**). Indeed, the expected gradual increase and decrease of FCGR3A (encoding CD16) and CD14, respectively, during CD14+ to CD16^+^ monocytes differentiation were not seen for scMerge and mnnCorrect data (**Fig. 5C-5F**), confirming an inferior selection by kBET. Given that monocytes from different technical batches should in theory distribute in the same pattern along the differentiation pseudotime, we exploited Kolmogorov-Smirnov (KS) test that measures pairwise consistency between distributions of different batches, with an average of three pairwise comparisons for overall consistency. We found RBET outcompeted LISI and kBET, because Combat selected by RBET resulted in the lowest KS value, which indicates the best consistency (**Fig. 5G**).

**Figure 5.**
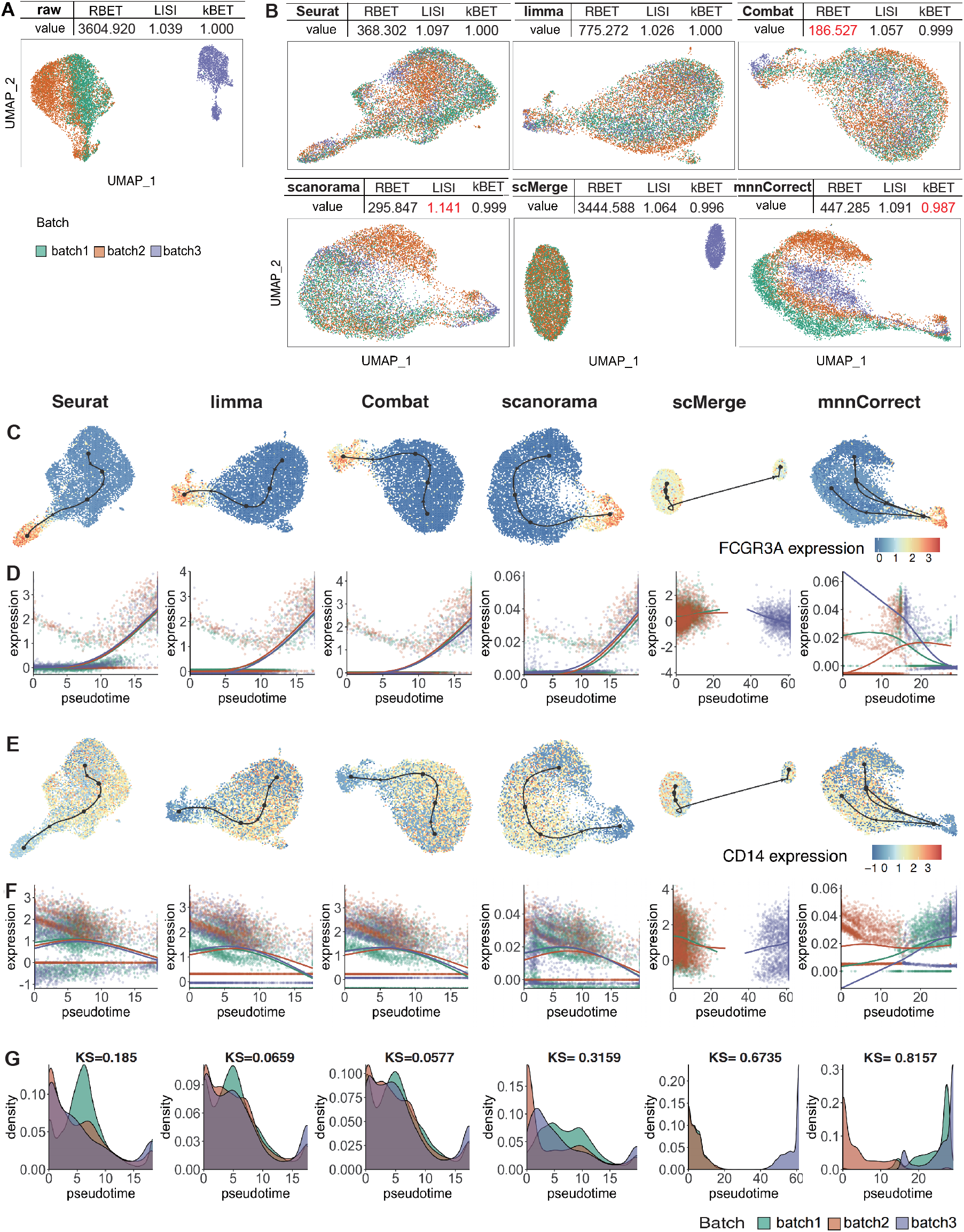
RBET correctly identifies the optimal BEC method in monocyte data. **(A)** UMAP visualization of original dataset colored by batches. **(B)** UMAP visualization of integrated data colored by batches. The table in the title of each sub-figure gives the BEC method used for the integration and the evaluation of integration result given by RBET, LISI and kBET, whose optimal choices are colored in red. **(C-D)** Expression profile of FCGR3A along the trajectory in UMAP (**C**) or along the peudotime (**D**). **(E-F)** Expression profile of CD14 along the trajectory in UMAP (**E**) or along the peudotime (**F**). (**G**) Distributions of cell numbers along the pseudotime in different batches. The mean value of KS statistics measures the consistency between these distributions. The legend for BEC tools in **(D-G)** is the same as in **(C)**.

Next, we challenged a multiple-lineage scenario with a complex CD8+ T cell dataset extracted from ovarian, lung and colorectal cancers, including 40 samples and 16,133 cells (**Fig. 6A, 6B**)^10^. We found that RBET and LISI preferred Seurat and mnnCorrect, respectively, but kBET gave a constant value of 1 to all BEC tools (**Fig. 6C**), suggesting a failure in evaluating dataset with high batch effect size as predicted by simulated data (**Fig. 2G**). This dataset consisted of two common CD8+ T cell lineages, including CD8+ Terma (cytotoxic) and CD8+ Tex (exhuastic), which are both differentiated from CD8+ Tn (naïve) cells as previously validated^10^. Only Seurat, limma, Combat and scanorama correctly identified the two CD8+ lineages using Slingshot, while lineages derived from scMerge and mnnCorrect all converged into a single end that is not consistent with CD8+ T cell biology (**Fig. 6C, S6A**). Unlike the monocyte dataset that contained purely technical batches, this T cell dataset was derived from different cancer types. Therefore, their cell distribution along the pseudotime trajectory may vary accordingly (**Fig. 6D**), but they should still more or less reflect the cell fraction characteristics of T cell subtype in certain cancer. For example, colorectal cancer had relatively less Temra cells, and ovarian cancer had less Tex cells (**Fig. 6A**), which was correctly reflected in Seurat, Combat and scanorama trajectories (**Fig. 6D**, note the last peak at the end of pseudotime for each lineage). We then searched for classical marker gene expression pattern along the lineage-dependent pseudotime of CD8+ T cell differentiation: (1) a gradual decrease of Tn marker CCR7 and a gradual upregulation of activation marker NKG7 along the trajectories; (2) the terminal stage marker genes CX3CR1, FGFBP2 for Terma and PDCD1, HAVCR2 for Tex, gradually increase along Temra and Tex lineages, respectively; (3) the Tem marker GZMK expresses similarly high in the middle of both trajectories, since Tem is a common progenitor state before differentiation into Temra and Tex stages. According to this expected gene expression pattern, trajectories derived from Seurat, limma, Combat and scanorama generated acceptable results (**Fig. 6E**), but RBET’s top choice Seurat outperformed other tools for this complex dataset in term of marker gene consistency and cell distribution consistency (**Fig. 6E, 6F**). Interestingly, CD8+ Tex lineage is the known target for anti-PD-1/PD-L1 treatment, and PDCD1 that encodes PD-1 protein expressed relatively higher in Tex lineage for lung cancer as compared to other cancers in Seurat processed trajectory (**Fig. S6B**), which correctly reflects current superior clinical outcome of anti-PD-1/PD-L1 treatment for lung cancer.

**Figure 6.**
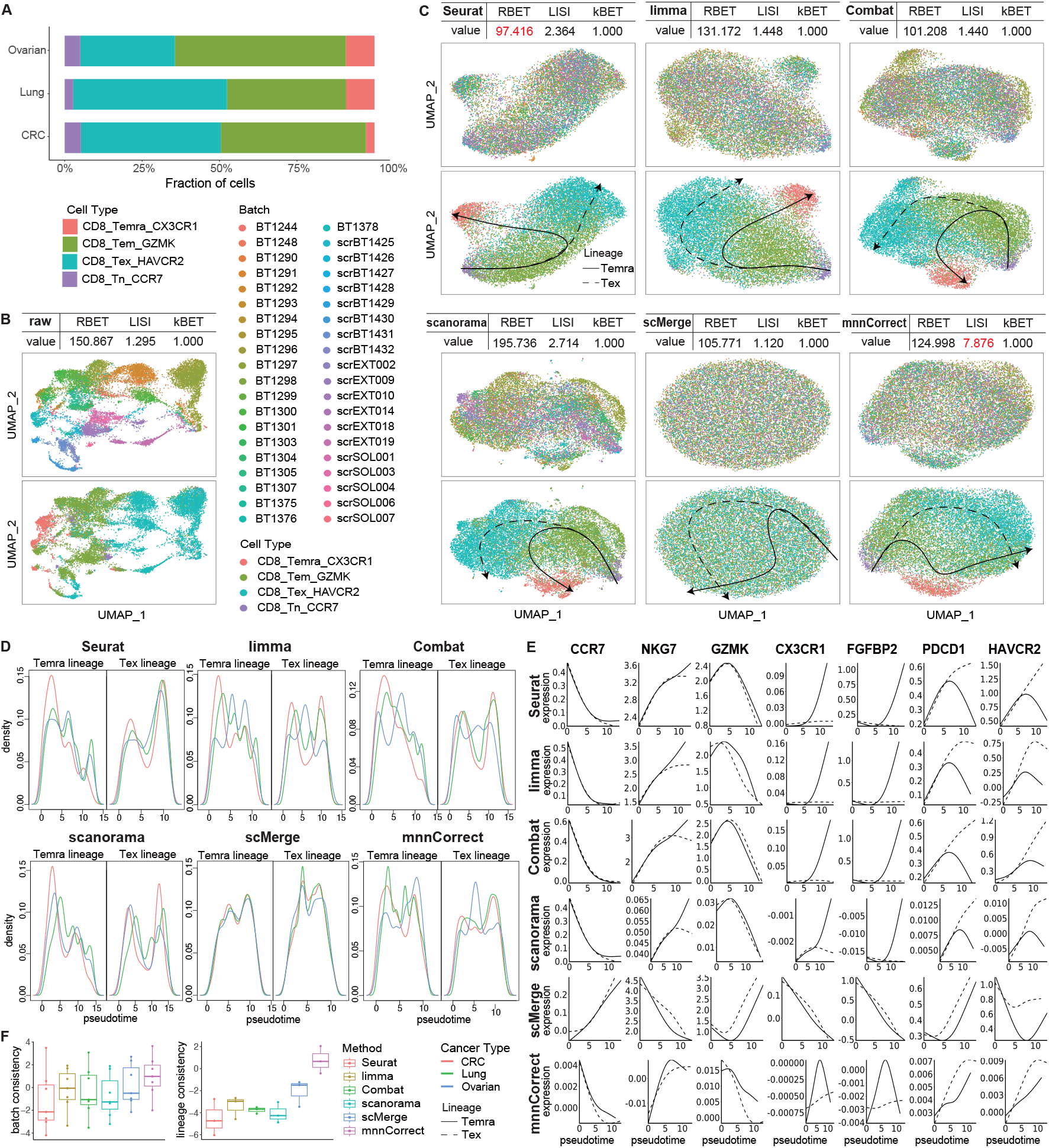
RBET chooses the optimal BEC method validated by trajectory analysis in CD8+ T cell dataset. **(A)** Histogram of cell type composition per cancer type. **(B)** UMAP visualization of original dataset colored by batches (up) and cell types (down). **(C)** UMAP visualization of integrated data colored by batches (up) and cell types (down), together with inferred trajectories (down). The table in the title of each sub-figure gives the BEC method used for integration and the evaluation of integration result given by RBET, LISI and kBET, whose optimal choices are colored in red. **(D)** Density plots for colorectal cancer (CRC), lung cancer and ovarian cancer along the two trajectories. See cancer type legend in **(F). (E)** The expression profile of 7 marker genes along the pseudotime, split by lineages, solid lines for Temra lineage and dashed lines for Tex lineage. **(F)** The consistency of the expression profile across batches for 7 marker genes (left). The consistency of the expression profile across lineages for 3 marker genes (CCR7, NKG7 and GZMK) (right).

### RBET facilitates cell-cell communication analysis

To investigate the influence of BEC on cell-cell communication analysis, we focused on the testicular data from healthy males^33,34^, which contained two batches and covered all the developmental stages from spermatogonial stem cells (SSCs) to mature spermatozoa. With almost identical composition of cell types, the batch influence was mainly in late stage (**Fig. 7A, 7B**). This batch effect was still retained after limma and Combat correction as their sperm cells were still split by batches (**Fig. 7C**), while other four methods gave acceptable batch correction on UMAP. To facilitate RBET, 12 housekeeping genes are curated from literatures and 4 of them are selected as RGs (**Table S4**). RBET, LISI and kBET selected scanorama, mnnCorrect and scMerge as methods of choice, respectively (**Fig.7C**). Data-based RBET gives the same selection as RBET (**Table S5**). To make further comparison, we dived into downstream analyses including cell type annotation, trajectory inference and cell-cell communication analysis. The scMerge result suffered from uniform marker gene expression across different stages and a discontinuous trajectory (**Fig. S7A, S7B**), hinting overcorrection, while expected marker gene expression patterns and developmental trajectory were only seen from scanorama, Seurat, and mnnCorrect data.

**Figure 7.**
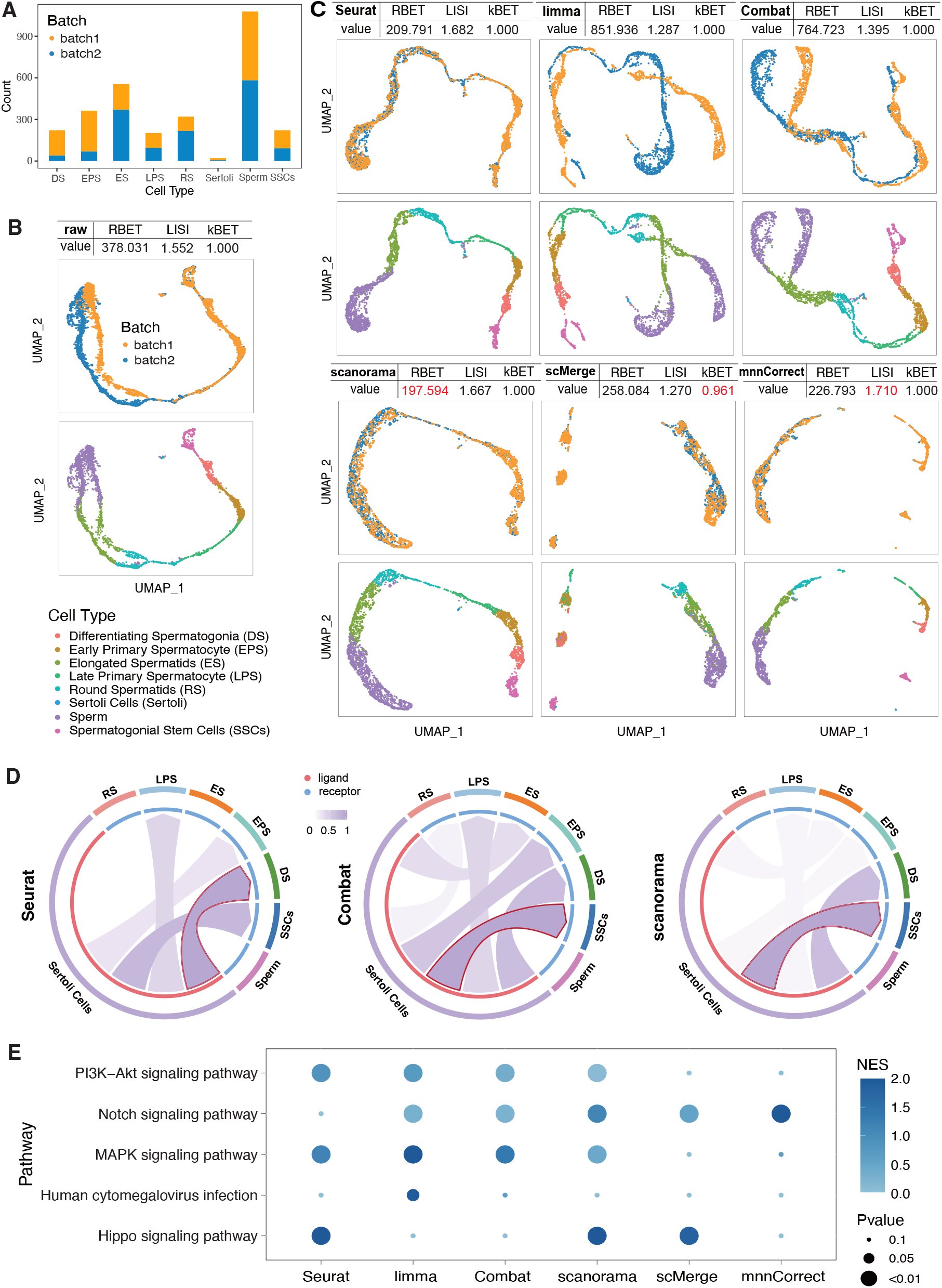
RBET achieves the best performance in testicular data with small batch effect. **(A)** Histogram of cell composition in each batch. **(B)** UMAP visualization of original dataset colored by batches (up) and cell types (down). **(C)** UMAP visualization of integrated data colored by batches (up) and cell types (down). The table in the title of each sub-figure gives the BEC method used for the integration and the evaluation of integration result given by RBET, LISI and kBET, whose optimal choices are colored in red. **(D)** Strength of communications from Sertoli cells to other germ cells. The arrow outlined in red indicates the strongest communication strength. **(E)** Pathway activity of five pathways reported to be enriched by intercellular signaling from Sertoli cells to SSCs.

Next, we focused on cell-cell communication from Sertoli cells to other cell types using CellChat package, since the former serves as germ cell ecological niches that support germ cell development^35^. Both Combat and scanorama gave the highest communication score to the Sertoli-SSC pair, which is consistent with previous report^33^. In contrast, Seurat wrongly assigned Sertoli-DS as the top hit. Further, we explored ligand-receptor (L-R) interactions and inferred their related KEGG pathway activity between cell types. For the four established Sertoli-SSC pathways that are crucial for spermatogenesis^36^, only scanorama recovered all of them (**Fig. 7E**). Human cytomegalovirus infection has gametotoxic effect on male germ cells and contributes to male infertility, at least partly through disruption of Sertoli-germ cell interaction^37^. Since this pathway should not be activated in the dataset derived from healthy donors, it can serve as a negative control. In this setting, data processed by limma wrongly identified this pathway, again highlighting the importance of choosing right BEC tools for downstream analysis. Furthermore, scanorama corrected data identified transcription factor (TF) SMAD1 and HES1 as key L-R downstream signaling effectors for Sertoli-SSC communication and spermatogenesis, which is consistent with previous findings that SMAD1/2 and HES1 are intracellular effectors of Notch signaling pathway for SSC proliferation and differentiation (**Fig. S7D**)^33,38^. However, the same analysis based on other BEC tools only identified one or none of these important TFs. In summary, RBET correctly selected scanorama as the best BEC tool, which was validated by downstream cell clustering, trajectory inference and cell-cell communication analyses for the testicular dataset.

## Discussion

Several BEC benchmark studies have comprehensively evaluated the data integration outcomes using up to 14 metrics. Different conclusions were drawn as some recommended all-weather top performers while other suggested a scenario-based guideline of choices. Our analyses favor the latter, since a simple linear regression model Combat was suitable for simple biological or technical replicates with limited cell type complexity (**Fig. 5**) and nonlinear Seurat CCA and scanorama performed better in high batch effect and complex cell type composition scenarios (**Fig. 4, 6, 7**). Nevertheless, the use of different metrics can lead to opposite conclusion, for example, a former top rated Harmony is discouraged to be used for complex dataset in a later study^8,9,39^. This discrepancy may stem from the use of different test datasets, as well as different multi-metric systems, which is another dilemma for common practitioners due to the lack of consensus. Different metrics only assess, mostly indirectly, limited aspects of BEC performance, and it appears tedious to perform a 14-metric benchmarking in routine practice, not to mention the expertise required to interpret and rank metrics for datasets of different research fields. In this regard, a simple metric, which takes full consideration of batch detection, data overcorrection, high batch effect robustness and scalability, is an unmet need.

These features are of particular importance for large scale and complex datasets that contain large batch effect, and for those applications that require full expression matrices to perform multiple downstream analyses. But the exisiting BEC evaluation algorithms, including LISI and kBET, are not fully armed. For example, they were not sensitive to overcorrection (**Fig. 3E**), and their batch detection capacity became quickly exhausted as the batch effect size increased (**Fig. 2G** for LISI and kBET in simulated data**; Fig. 6C, 7C** for kBET in real data).These may result in sub-optimal BEC choice and lead to wrong biological conclusions in downstream analyses. Previous benchmarking studies largely focused on metrics for cell identity annotation, with only a few downstream analyses including trajectory conservation^8,9^. We went one step further to explore single- and multi-lineage trajectories according to known marker expression patterns along the differentiation pseudotime, and examined their expression consistency, as well as cell distribution consistency across batches. Moreover, we performed cell-cell communication analysis for the testicular dataset and identified known interaction paired as well as key L-R downstream signal pathways and TFs (**Fig. 7**), where RBET demonstrated its superiority.

The idea of RBET is inspired by the concept of housekeeping genes or RGs, which are stably and abundantly expressed with moderate expression variation across cells. This reminds us that an ideal BEC method should achieve the similar RG expression profile across batches without losing gene expression variation. The RBET workflow has only two simple steps including the RG matrices construction and MAC test on the two-dimensional vector space, and a lower RBET score indicates better performance of BEC. Different strategies for constructing RG matrices, i.e., literature-based and data-based RGs, give consistent decisions in our examples (**Fig. 4B, 5B, 6B, 7B, Table S5**). In summary, for both simulated and multi-scenario real datasets, RBET outperformed LISI and kBET in terms of detecting power, type I error control, overcorrection awareness, computational efficiency and large batch effect robustness.

Finally, previous benchmark works are highly appreciated for providing a multi-scenario guideline for choosing optimal BEC tools, but this approach also have limitations: 1) re-evaluation required when new BEC tools emerge, 2) not always easy to categorize the dataset before analysis, and 3) all comparisons are performed using default BEC parameters while changing these represents another level of BEC optimization. As demonstrated in **Fig. 2**, changing the k.anchor value to 10 for Seurat CCA gave the lowest RBET value thus an optimal choice. Given the multi-feature and easy to use advantages of RBET, we propose to apply RBET for routine BEC optimization practice, either by comparing different BEC tools or by finetuning parameters for a favorite BEC tool. And this may open an era for case-specific optimal decision making for BEC tools instead of following a general guideline. Last but not least, given the concept of RGs and dimensionality reduction that can be similarily applied to other single-cell modalities, i.e. scATAC-seq, CITE-seq and MERFISH, it is conceivable to expand the ultilty of RBET for broader BEC tasks.

RBET is provided as an R package with curated lists of housekeeping genes for multiple tissue types, RG calculation tool and MAC test function. In addition, the package offers user friendly interface to common BEC tools like Seurat, which is available at https://github.com/zlyx26/RBET.

## Methods

### Literature-based reference genes selection

Literature-based reference genes are selected from experimentally validated housekeeping genes specific to the tissue or cell type of interest. It goes in four steps as follows. **(Step 1)** Annotate cell types in each batch by manual annotation or automatic annotation tools like ScType^26^. **(Step 2)** Select experimentally validated housekeeping genes specific to the current tissue or cell type as candidates from existing literature. To facilitate the analysis, we provide a database of experimentally validated housekeeping genes for 23 tissues or cell types collected from literature (Table S6), also available in our R package. **(Step 3)** Rank the candidate genes according to the number of cell types where they are differentially expressed across batches. First, the scRNA-seq data is log-transformed by log(1+count). Next, we use Mood’s median test to test whether a candidate gene is differentially expressed in each cell type (mood.medtest function in R package *RVAideMemoire*, v0.9-83), and get the p-value *p*_*ij*_ for the *i*-th candidate in the *j*-th cell type. Then, we count the number *n*_*i*_ of cell types where the *i*-th gene is differentially expressed across batches, i.e., *n*_*i*_ = ∑_*j*_ *I,p*_*ij*_ < 0.050. Finally, we rank the candidate genes according to *n*_*i*_. **(Step 4)** Those candidates with the largest *n*_*i*_ are taken as reference genes, and each batch should have at least 1 RG. Note that the number of reference genes should be at least 2.

### Data-based reference genes selection

In case of no reported tissue-specific housekeeping genes or difficult cell annotation, data-based reference genes selection is also provided, which chooses reference genes directly from data. Similar with literature-based method, it selects genes that express invariably in each batch and differentially across batches. It contains the following five steps. **(Step 1)** Cluster cells in each batch (FindClusters function in R package *Seurat*, v4.1.0). **(Step 2)** Select genes that express invariably in clusters. For each batch, we subset the expression of the *i*-th gene in the *j*-th cluster as *e*_*ij*_, and compute an index as the standard deviation of *e*_*ij*_ divided by the mean of *e*_*ij*_, that is, *I*_*ij*_ = *sd*(*e*_*ij*_)/mean(*e*_*ij*_). Then, we rank the genes according to the index from small to large in each cluster, and count the number of clusters *c*_*i*_ where the *i*-th gene is in the top *r*%. Next, we rank the genes according to *c*_*i*_ from large to small in each batch, and count the number of batches *b*_*i*_ where the *i*-th gene is in the top *r*%. Finally, we rank the genes according to *b*_*i*_ from large to small, and the genes in the top *r*% are taken as candidates in clusters. **(Step 3)** Select genes that express invariably across clusters. For each batch, the expression of the *i*-th gene is taken as *e*_*i*_ and the mean expression of the *i*-th gene in each cluster is taken as *m*_*i*_. We compute an index *I*_*i*_ = *sd*(*m*_*i*_)/mean(*e*_*i*_). Then we rank the genes according to the index from small to large in each batch, and count the number of batches *b*_*i*_ where the *i*-th gene is in the top *r*%. Finally, we rank the genes according to *b*_*i*_ from large to small, and the genes in the top *r*% are taken as candidates across clusters. **(Step 4)** We first take an intersection of candidates in clusters and across clusters. Then, we count the times *n*_*i*_ when the *i*-th candidate gene expresses differentially across batches by comparing pairwise clusters in pairwise batches (mood.medtest function in R package *RVAideMemoire*, v0.9-83), and rank the candidates according to *n*_*i*_ from large to small. **(Step 5)** The top *l* candidates are taken as reference genes. Note that the number of reference genes should be at least 2. Our method is quite stable under different *r* and *l*, and here the default values are set as *r* = 5 and *l* = 30.

### RBET statistic for batch effect detection

#### Concept of RBET statistic

Let **X** ∈ ℝ^***n*×*p***^ and **Y** ∈ ℝ^***m*×*p***^ be the gene expression matrices of two different batches, where *n* and *m* are cell numbers and *p* is the gene number. Input matrices with different genes can be inner joined together by shared genes, and thus the gene numbers become the same. We notice that samples from two different batches can be well regarded as samples from two different distributions. In such a view, the problem of batch effect detection naturally fits into the framework of two-sample distribution test, with the null hypothesis that **X** and **Y** have the same distribution:

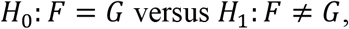

where *F* and *G* are distribution functions of **X** and **Y**, respectively.

For the case *p* = 1, this problem is well studied in statistical literature^15^. However, only few methods have been proposed for the high-dimensional cases. Recently, we introduced a class of MAC statistics for this problem^16^. Theoretical analysis has demonstrated that MAC has an upper bound of order (log(*nm*)) when there is no batch effect, which is much smaller than the lower bound of order (*n* + *m*) when batch effect is present^16^. Notably, MAC outperforms other methods in cases *p* < 10, but it is almost inapplicable when *p* exceeds 10. Hence, due to the substantial size of genes in our datasets, i.e., *p* could be as large as thousands, MAC cannot be directly employed.

Motivated by the fact that dimensionality reduction tools such as tSNE and UMAP tend to preserve the neighborhood characteristics of each point in the original space, we propose to use them to map the original dataset into a two-dimensional vector space, and then MAC statistics can be applied. Specifically, we put **X** and **Y** together to form a large (*n* + *m*) **×** *p* matrix, and reduce it into a two-dimensional observation **Z** ∈ ℝ^(*n*+*m*)**×**2^. **Z** can be partitioned into 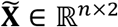 and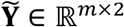, corresponding to **X** and **Y**, respectively. In this way, we obtain a two-dimensional vector for each batch in the same vector space which well-preserves the original neighborhood characteristics. Next, we implement MAC with *p* = 2 to detect the batch effect among 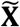 and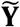, namely that among **X** and **Y**.

Let ***x*** = {***x***_***i***_: *i* = 1, 2, …, *n*} be the collection of *n* rows of 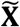, and ***y*** = {***y***_***i***_: *i* = 1, 2, …, *m*} be the collection of *m* rows of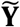. Thus, ***x*** and ***y*** are mapped samples from two different batches. Let (***x***_***i***_, ***y***_***j***_) be any pair of mapped samples, where ***x***_***i***_ = (*z*_*i*,1_, *z*_*i*,2_) and ***y***_***j***_ = (*z*_*j*,1_, *z*_*j*,2_), and 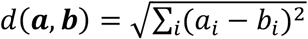 be the Euclidean distance between two vectors. Define

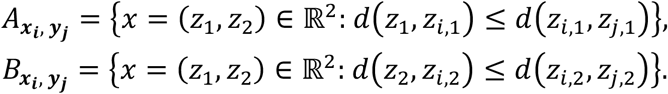

Then, we divide the whole sample space ℝ^2^ into four disjoint parts,

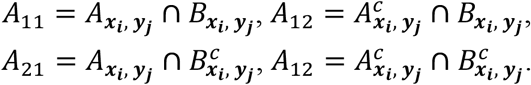

Let 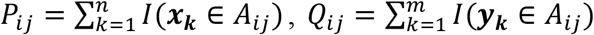, *R*_*ij*_ = *P*_*ij*_ + *Q*_*ij*_, *i, j* = 1, 2, then the local statistic *T*_*ij*_ that compares the distribution of ***x***_***i***_ and ***y***_***j***_ is defined as

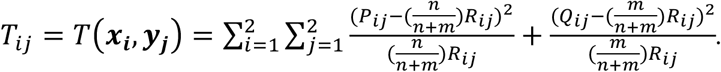

It should be noted that if any *R*_*ij*_ equals to zero, *T*_*ij*_ is set to zero. Finally, MAC statistic takes the maximum of all the local statistics *T*_*ij*_, i.e.,

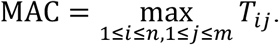

#### Sub-sampling strategy to improve computational efficiency

Since scRNA data often comprise hundreds or even thousands of cells, the calculation of MAC statistics on the whole sample is computationally expensive. Considering that similar cells have similar gene expression profiles and cluster together in the reduced vector space, we propose a sub-sampling strategy to improve the computational efficiency. To be more specific, we rank rows of ***Z*** according to its first column, and select samples whose row numbers equal to Δ **×** *j* with *j* = 1, 2, …, *k* = (*n* + *m*)/Δ, where Δ is the sampling gap. Then we calculate RBET based on the *k* selected samples. It should be noted that this sampling strategy simply reduces the number of local statistics used for defining MAC, without changing its theoretical properties.

### Compared methods for batch effect detection

RBET is compared with two other methods. (1) kBET^9^. It assumes that if there is no batch effect, the distribution of batch labels within any given neighborhood of equal size should mirror that of the entire dataset. Based on this idea, kBET applies Pearson chi-squared test to randomly selected neighborhoods of a fixed size, and then returns an overall rejection rate which reflects the degree of mixing in the dataset. A low rejection rate indicates that the two datasets are well mixed and there is no or small batch effect. R package *kBET* (v0.99.6) is used here and all parameters are set by default. (2) LISI^7^ based on Inverse Simpson Index (ISI). LISI first assigns weights to adjacent cells by constructing Gaussian kernel-based distributions of neighborhoods, and then calculates ISI as the effective number of batches in each cell’s neighborhood. The mean of LISI from all the neighborhoods is defined as the overall number of batches. R package *lisi* (v1.0) is used for calculation with default parameters.

### Simulation

#### Gaussian examples

In these examples, we generated synthetic data following Gaussian distributions. We assumed that *X* followed a two-dimensional Gaussian distribution, *F* = *N*((0,0), *I*), and considered three different cases for the distribution of *Y, G*. (G1) *Y* followed *N*((*μ, μ*), *I*) with *µ* varying from 0 to 1; (G2) *Y* followed *N*((0,0), *σI*) with *σ* varying from 1 to 4; (G3) *Y* followed *N*((0, 0), *Σ*), where *Σ* = (*σ*_*ij*_), *σ*_11_ = *σ*_22_ = 1, and *σ*_12_ = *σ*_21_ = *ρ* with *ρ* varying from 0 to 1. For each case, we sampled *n* (= 50, 100, 200) independently and identically distributed observations from *F* and *G*, respectively.

#### Examples with simulated gene expression data

In these examples, we applied R package *splatter* (v1.18.2)^40^ to simulate gene expression counts for two different batches, each with 2000 genes and 400 cells. Then, we tested the power of three batch effect detection tools (RBET, LISI and kBET) across different numbers of cell types, batch effect sizes, and the proportions of cells with batch effect. The batch effect sizes ranged from 0 to 0.05.

First, we considered the parameter *k* in the sub-sampling strategy of RBET. In this case, all the cells were simulated to have batch effect, and the number of cell types changed from 1 to 4.

Second, we compared the power of different methods under two different settings: (S1) all the cells had batch effect; (S2) only a proportion of cells had batch effect. (S2) was further divided into 3 cases with different numbers of cell types in each batch. (S2a) There were two cell types in each batch. The batch effect only occurred in the first cell type, and the proportion of cells in it varied from 20% to 80%. (S2b) There were 4 cell types in each batch, and 60% of cells had batch effect. Cells with batch effect were uniformly distributed in *n* cell types, with *n* ranging from 1 to 3. For instance, if we set *n* = 3, the cell numbers in these three cell types were all 400 **×** 60%/3 = 80, and the cell number in the last cell type was 400 **×** (1 − 60%) = 160. (S2c) There were 8 cell types in each batch, and 60% of cells had batch effect. Cells with batch effect were uniformly distributed in *n* cell types, with *n* ranging from 1 to 7.

#### Value variation

We studied the impact of batch effect size on the values of three batch effect detection tools. For a given effect size, we simulated *m* batches of data, each with 2000 genes and 400 cells, and *m* varied from 2 to 10. The batch effect sizes ranged from 0.1 to 1.

#### Overcorrection problem

We also simulated data for the overcorrection problem, which contained two batches, each with 4 groups. All the cells had batch effect, with the effect size fixed to 0.5. The data were corrected by Seurat (v4.1.0 or v3.2.0) under different k.anchor. HVGs were identified in each batch before and after batch correction, using FindVariableFeatures(selection.method = “mean.var.plot”) function in R package *Seurat*. Noted that R package *splatter* (v1.18.2) outputted a differential expression factor for each gene, DEFac, we took all the genes with DEFac unequal to 1 as true DEGs. DEGs after batch correction were identified by FindAllMarkers function in R package *Seurat*.

#### Running time

We compared the total running time of different methods for 200 repetitions in datasets with two batches but different sizes. (a) 1000 cells and 2000 genes in each batch; (b) 2000 cells and 4000 genes in each batch; (c) 4000 cells and 8000 genes in each batch.

### Analysis details

#### Data processing

The raw data and the integrated data were mainly analyzed with Seurat (R package, v4.1.0)^17^. After data normalization and scaling, the top 2,000 variable genes were calculated by FindVariableFeatures function and used to construct principal components (PCs). PCs covering the highest variance were selected according to the elbow and Jackstraw plots. Clusters were calculated by FindClusters function and visualized using the Uniform Manifold Approximation and Projection for Dimension Reduction (UMAP).

#### Tools for integrating scRNA-seq datasets

Most downstream analysis, such as differential gene expression analysis and gene set enrichment analysis, require expression matrices as input. Here, we aimed to evaluate different BEC methods based on the validity and accuracy of subsequent downstream analysis. Therefore, six commonly used BEC tools that could return gene expression matrices with full dimensionality were applied for data integration: Seurat (R package, v4.1.0), limma (R package, v3.50.3), Combat (R package, v3.42.0), scanorama (R package, vl.7.2), scMerge (R package, vl.70.0), and mnnCorrect (R package, v1.12.3).

#### Cell annotation

Annotation of cell clusters is a fundamental step for single-cell analysis. In this study, we utilized the marker-based automatic annotation tool ScType^26^ together with the marker genes specific to each tissue or cell type. ScType matched the gene expression patterns of data to the known cell types in order to label individual cells or cell clusters^26^. We compared the annotation results with real cell tags under three metrics, which were accuracy (ACC), adjusted rank index (ARI; R package *aricode*, v1.0.2), and normalized mutual information (NMI; R package *aricode*, v1.0.2). Additionally, we evaluated the quality of annotated clusters using silhouette coefficient (SC; R package *cluster*, v2.1.6), which quantified how similar a cell was to its own cluster compared to other clusters.

#### Trajectory inference

Trajectory analysis aims to construct an ordered sequence of cells to describe the dynamic changes in single-cell gene expression levels. Here, Slingshot (R package *slingshot*, v2.2.1) was applied to perform pseudotime analysis on the integrated datasets, which was recommended by previous studies^28^. It took the low dimensional embedding of gene expression and a vector of cluster labels as the input. Here, the labels were predicted using k-means clustering method (kmeans function in R package *stats*, v4.1.1). Then, we utilized getLineages function in *slingshot* to fit a minimum spanning tree (MST) to identify lineage. In accordance with established biological knowledge, we set the root for CD8+ T cell dataset as CD8_Tn_CCR7 and the root for testis dataset as SSCs. However, monocyte dataset lacked explicit cell type annotation, and thus the trajectory was inferred automatically without a prespecified root. The final output was stored in a SlingshotDataSet object containing inferred lineage information.

#### Measuring the consistency of gene expression profiles along the pseudotime

The expression profile *e* of gene *G* could be fitted as a smoothing function of the inferred pseudotime *t*, denoted as *e*(*t*) (smooth.spline function in R package *stats*, v4.1.1). Assuming that *e*_(_(*t*) and *e*.(*t*) were the fitted curves of expression profiles of gene *G* in two different batches or lineages, we defined the consistency between these two curves as the averaged distance between them, i.e.,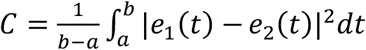. It was actually the mean of |*e*_1_(*t*) – *e*_2_(*t*)|^2^ by taking *t* as a uniformly distributed random variable on [*a, b*]. This provided us a simple measurement by first sampling *n* instances, {*t*_1_, …, *t*_*n*_}, from *U*[*a, b*], and then estimating the averaged distance as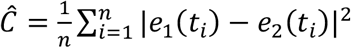.

In this paper, we took *n* = 1000, which was large enough to get an accurate estimation. To mitigate the effect of gene expression size, we normalized 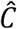 by the square of the mean expression (*μ*_*G*_), i.e.,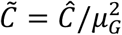. In case of multiple batches, we calculated their pairwise consistency and took the average.

#### Cell communication analysis

The biological behavior of cells is regulated by the intercellular and intracellular signaling network, which is known as cellular communication. Here, we performed cell-cell communication analysis on the integrated datasets using CellCall (R package *cellcall*, v1.0.7)^33^. First, we inferred the overall communication scores between different cell types using TransCommuProfile function. Next, we identified crucial pathways involved in communication by performing a pathway activity analysis with getHyperPathway function. Finally, we visualized the intercellular signaling from one cell type to another in sankey plot (LR2TF and sankey_graph functions).

## Data availability

All the datasets used for analyses in this work are publicly available online, deposited in the NCBI’s Gene Expression Omnibus database and ArrayExpress. In particular, we used a pancreas dataset^24^ containing three batches of single-cell expression data obtained by different sequencing technologies, CelSeq (GSE81076)^41^, CelSeq2 (GSE85241)^27^ and SMART-Seq2 (E-MTAB-5061)^42^, a human monocyte dataset^30^ under accession code GSE146974, a CD8+ T cell dataset extracted from Qian et al.^10^ under accession number E-MTAB-8107, E-MTAB-6149 and E-MTAB-6653, taking each sample as a batch, and a testis dataset^43^ of testicular transcriptome sequencing data from healthy males under accession code GSE112013.

## Code availability

The code for RBET is written in R and Rcpp, and can be obtained at https://github.com/zlyx26/RBET.

